# Phenotype and gene ontology enrichment as guides for disease modeling in *C. elegans*

**DOI:** 10.1101/106369

**Authors:** David Angeles-Albores, Raymond YN Lee, Juancarlos Chan, Paul W Sternberg

## Abstract

Genome-wide experiments have the capacity to generate massive amounts of unbiased data about an organism. In order to interpret this data, dimensionality reduction techniques are required. One approach is to annotate genes using controlled languages and to test experimental datasets for term enrichment using probabilistic methods. Although gene, phenotype and anatomy ontologies exist for *C. elegans*, no unified software offers enrichment analyses of all the ontologies using the same methodology. Here, we present the WormBase Enrichment Suite, which offers users the ability to test all nematode ontologies simultaneously. We show that the WormBase Enrichment Suite provides valuable insight into different biological problems. Briefly, we show that phenotype enrichment analysis (PEA) can help researchers identify disease phenologs, phenotypes that are homologous across species, which can inform disease modeling in *C. elegans*. The WormBase Enrichment Suite analysis can also shed light on RNA-seq datasets by showing what molecular functions are enriched, which phenotypes these functions are implicated in and what tissues are overrepresented in the dataset. Finally, we explore the phenotype-anatomy relationship, showing that a small subset of highly specific tissues are disproportionately likely to cause an Egl phenotype, but inferring tissue expression from an Egl phenotype is limited to the largest tissues.

## Introduction

The last decade has seen an explosion of techniques capable of genome-wide measurements. Some examples of genome-wide tools include RNA-seq [1] to measure gene expression or CHIP-seq [2] to measure protein binding to chromatin. These tools are capable of generating large quantities of data. Understanding these data, and generating hypotheses from them remains challenging. A common approach used to understand these datasets is to reduce the dimensionality of the data via enrichment analyses of ontologies [3], which helps researchers understand what terms are overrepresented beyond random levels. By analyzing overrepresented terms in aggregate, researchers can better understand what biological processes were most affected in a given experiment, and form hypotheses about what is happening [4]. This approach is limited by what ontologies can be tested for enrichment. The best-known ontology for biological research is the Gene Ontology (GO), which provides a controlled language to describe molecular and cellular functions of genes [3]. In *C. elegans*, gene, tissue and phenotype ontologies exist with which to describe *C. elegans* anatomy and phenotypes respectively [5, 6]. These ontologies are curated by professional curators at WormBase, which is a repository of all *C. elegans* data [7]. However, enrichment tools only exist for gene and tissue ontologies in the community today (see for example [8–10]). Another limitation is that tissue enrichment testing is not offered on the same websites as GO enrichment testing, which requires users to test their data on different websites that may or may not use different methodologies to detect enrichment.

Another way to use enrichment tools is for evolutionary comparison purposes. In molecular biology it is often useful to know when a gene is homologous between two species—that is to say, common by descent—because knowledge of homology often brings with it knowledge of function. Indeed, many important gene regulatory networks (GRNs) are conserved between organisms as highly diverged as nematodes and humans (for example, see [11]). While genes and GRNs may be conserved between species, their outputs often differ. For example, the gene Pax6 (Eyeless in *Drosophila melanogaster*) is involved in eye formation in humans and fruit-flies [12]. Although nematodes have conserved this gene, they do not have eyes [13, 14]. The concept of a phenolog has been put forward to explain relationships between phenotypes that have the same underlying genetic regulatory network [15, 16]. Formally, two phenotypes are phenologs of each other if the orthologs of the genes that cause a phenotype in an organism cause a second phenotype in another.

To study a clinically relevant disease in a non-human, an appropriate model has to be established. A straightforward method towards establishing a disease model in *C. elegans* is to link a disease to a causal gene, then to identify the homologous gene in *C. elegans* and then to study the function of the genetic homolog to extrapolate back to humans. However, this method relies on the existence of known disease genes and requires that the homolog have a phenotype that can be reliably identified and studied. A fundamentally different way to establish a disease model in *C. elegans* would be to identify the phenologs of the disease to be studied in *C. elegans* by identifying disease-associated human genes in an unbiased manner through genome-wide association studies (GWAS) and identified candidate homolog genes in *C. elegans*. The orthologs can be used to identify *C. elegans* disease phenologs, which can in turn be used as the basis for screens to identify genes that are associated with that phenolog. Approaches similar to this have been successfully used in the past to make non-obvious links between phenotypes in different species [15].

The concept of a phenolog can also be useful when applied within a species. In *C. elegans*, not all phenotypes are equally easy to study. Although genome-wide measurements can help elucidate the genetic network underlying a phenotype, devising screens to test which genes are functionally important can be difficult. A common strategy to study phenotypes that are difficult to screen is to select an easier-to-screen phenolog, and to test positive hits for the true phenotype of interest afterwards. For example, candidate genes to extend the *C. elegans* lifespan can be first screened for using heat shock survival genes involved in *C. elegans* aging sensitivity can be identified using stress assays [17, 18]. Currently, selection of screening phenotypes is performed based on researcher experience. By formalizing phenotype enrichment analysis as a tool with which to analyze gene sets, researchers should be able to formally establish phenologs, which has consequences for screen design.

An additional problem with genome-wide queries of *C. elegans* states (be they developmental, such as L1, L2, dauer; behavioral states such as awake versus asleep; or other) is that they do not always have a straightforward interpretation in terms of phenotypes. In these situations, researchers must rely on intuition to select a phenotype for which to screen. As a result, many hits may go unexplored that would prove fruitful. The question of how to design a screen that is maximally informative is an important question that has so far not been addressed within this community.

To facilitate understanding of large datasets, and to make discovery of phenologs easier, we have completed an enrichment tool suite in WormBase that allows users to rapidly perform phenotype, tissue and gene ontology enrichment analyses (PEA, TEA and GEA respectively) on curated *C. elegans* ontologies using the same methodology for each one. We applied our tools towards the unbiased discovery of phenologues of multigenic, complex diseases including systemic lupus erythematosus, obesity and obesity-related traits, and rheumatoid arthritis by using genes associated with these diseases via genome-wide association studies. We illustrate the utility of the complete enrichment suite for finding new relationships in complex data by analyzing a ciliary neuron transcriptome [19]. Finally, we show that the dictionaries generated for these enrichment analyses can help elucidate the contributions of specific tissues to specific phenotypes.

## Methods

### Implementing the enrichment analyses

All scripts were implemented in Python 3.5 [20]. We used pandas [21] and scipy [22] to write the statistical testing framework. Matplotlib [23] and Seaborn [24] libraries are used to generate all plots. Testing was performed using the WormBase version WS256. The WormBase Enrichment suite can be installed using pip via the command:

~~~
pip install tissue_enrichment_analysis
~~~

### Human disease phenolog identification

We used the GWAS EBI-NHGRI catalog [25] to extract information on all genome-wide association studies deposited there. We only selected traits that had > 300 associated genes. We identified 24 traits that met our criteria. Next, we used DIOPT [26] to identify candidate orthologs for the genes associated with these traits. Briefly, DIOPT combines a large number of methods for identifying orthologs and returns homolog candidates associated with a compound score. Depending on the score, orthologs can be considered ‘high’, ‘moderate’ or ‘low’ rank, reflecting confidence in the homology. Many-to-one and one-to-many homology relationships are allowed in DIOPT, reflecting a mixture of uncertainty and family expansion/reduction. For our study, we only accepted homolog candidates with ‘high’ or ‘moderate’ scores and we did not insist on a one-to-one relationship between genes.

After we identified worm orthologs for each trait, we reassessed how many traits still had > 100 gene candidates, and dropped all traits that had less than this for our analysis. We identified 18 traits that met this criteria. The gene lists for each of these 18 traits were then analyzed for gene, tissue and phenotype enrichment (see Fig.1). Tissue enrichment was performed using the WormBase Tissue Enrichment Analysis (TEA) tool [10].

**Figure 1.**
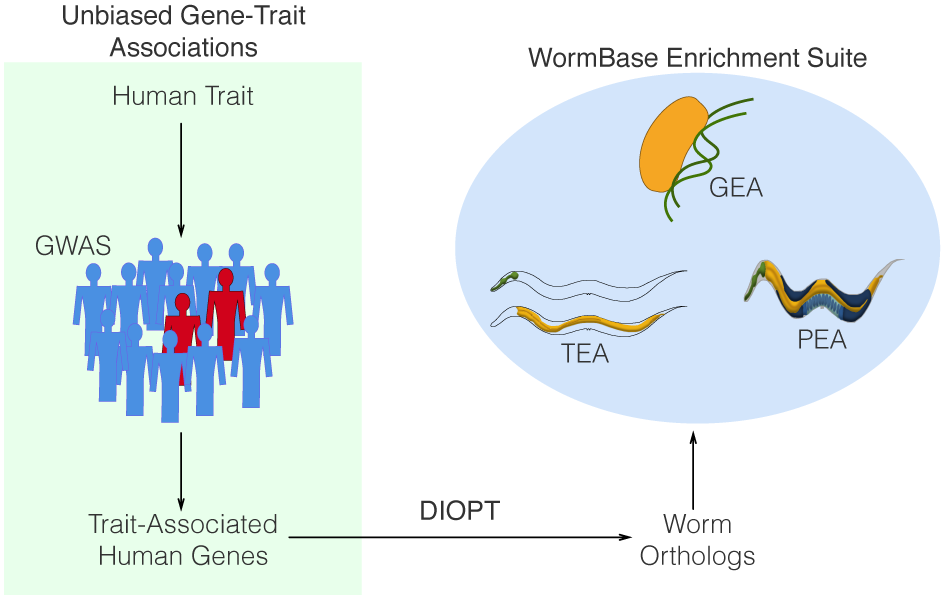
Experimental design for human-nematode phenologue identification. We used GWAS candidates from the EBI-NHGRI catalog to identify disease associated genes, then used DIOPT to identify candidate orthologs in *C. elegans*. Orthologs were used to run phenotype, gene and tissue ontology enrichment analyses to identify disease phenologues.

## Results

### Developing the WormBase enrichment suite

We developed the dictionaries for PEA and GEA using the same procedure as was used for TEA [10]. We generated a dictionary that included terms with at least 50 annotating genes or more and had a similarity threshold of 0.95 for PEA (the total number of terms in the dictionary was 251, annotated by 9,169 genes for the version WS256); and we generated a dictionary that included terms with at least 50 annotating genes or more and had a similarity threshold of 0.95 for GEA (the total number of terms in this dictionary is 271, annotated by 14,636 genes for the version WS256). Next, we benchmarked the dictionaries on the same gene sets as TEA and obtained enrichment of all the expected categories [27–34]. For example, on a gene set enriched for embryonic muscle genes [30], the top two enriched phenotype terms by q-value were ‘muscle system morphology variant’ and ‘body wall muscle thick filament variant’; the top two enriched GO terms were ‘myofibril’ and ‘striated muscle dense body’. For all the benchmarking results, see supplementary information. Having generated and validated our dictionaries, we proceeded to identify phenologs for several common human diseases.

### Applying the WormBase enrichment suite

To discover phenologs, we first needed to identify genes that contribute to a disease in an unbiased manner. One way to discover gene associations in an unbiased manner is to perform GWSA in human populations. Therefore, we used the GWAS NHGRI-EBI Catalog [25] to identify genes associated with human diseases. We found the best nematode candidate orthologs for these genes using DIOPT [26] and applied our enrichment suite to each of these gene regulatory networks.

### Systemic lupus erythematosus

Systemic lupus is an autoimmune disease that is believed to be polygenic in nature [35]. It mainly affects women and is characterized by painful and swollen joints, hair loss, and fatigue [36]. Since worms do not have a cellular immune system, we were interested in what phenologs corresponded to this disorder in *C. elegans*. To establish phenolog candidates, we obtained 283 genes associated with the disease via GWAS studies, and found 135 homolog candidates in *C. elegans*.

Lupus-associated orthologs were reasonably well annotated. Slightly more than half of the genes had at least one phenotype annotation (76/135) and almost all genes were annotated to at least one tissue or gene ontology term (104/135 and 115/135 genes respectively). We found that Lupus-associated orthologs were enriched in ‘aneuploidy’ (7 genes, *q* < 10^−1^) and ‘meiotic chromosome segregation’ (8 genes, *q* < 10^−1^). ‘Cell fate transformation’ (6 genes, *q* < 10^−1^), and ‘excess intestinal cells’ (5 genes, *q* < 10^−1^) were also overrepresented, as was ‘male tail morphology’ (6 genes, *q* < 10^−1^). Finally, the phenotype ‘nonsense mRNA accumulation’ was also enriched (5 genes, *q* < 10^−1^) (see Fig. 2).

**Figure 2.**
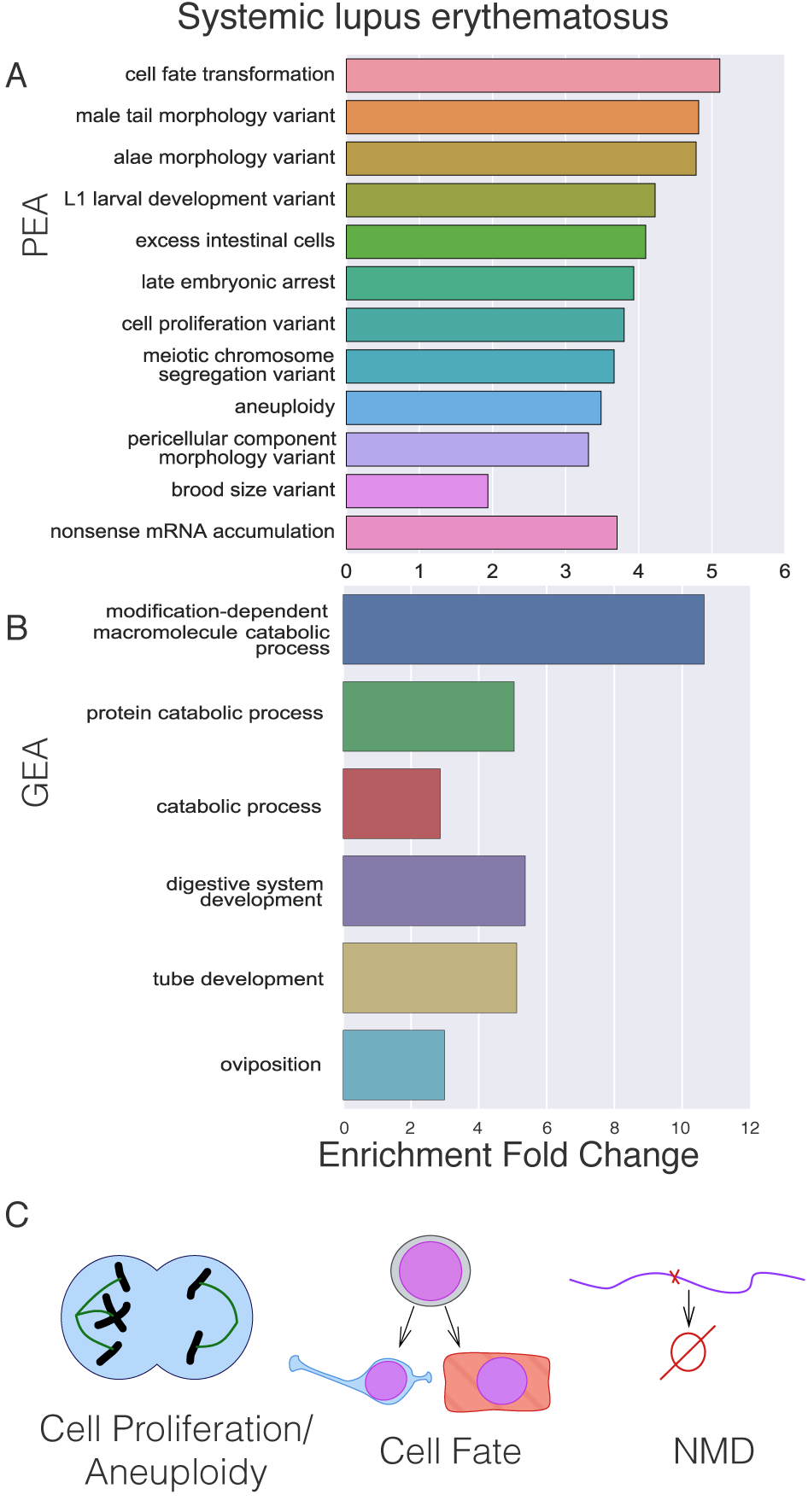
Phenologue identification for systemic lupus erythematosus. **A** Phenotype Enrichment Analysis. **B** GO Enrichment Analysis. **C** Lupus in *C. elegans* may be best represented by a combination of three phenotypes: Cell proliferation possibly accompanied by aneuploidy; cell fate transformations that may lead to dysmorphias; and a molecular phenotype involving impairment of the Nonsense-Mediated Decay (NMD) pathway.

TEA suggested that the ‘excretory duct cell’ (5 genes, *q* < 10^−2^) and the ‘posterior gonad arm’ are overrepresented in this dataset. We also found that the Pn.p cells P3.p through P8.p were enriched in this dataset (5 genes, *q* < 10^−1^). GO enrichment pointed at ‘modification-dependent macromolecule catabolic process’ (23 genes, *q* < 10^−15^) as a molecular function that characterizes this dataset. However, this GO term was enriched only due to a single gene family, the *skr* gene family. Almost the entire *skr* family was considered a candidate homolog to the SKP1 human gene, making the GO enrichment suspect.

Enrichment of the terms for ‘aneuploidy’, ‘meiotic chromosome segregation’, and ‘excess intestinal cells’ were largely driven by the same gene group, which includes *cki-1*, and several *skr* genes. On the other hand, ‘cell fate transformation’ and ‘male tail morphology’ reflected the involvement of developmental genes *let-23*, and *lin-12* among others. The term ‘nonsense mRNA accumulation’ was the result of *pept-3*, *smg-7*, *tsr-1*, *dhcr-7* and *F08B4.7*. Therefore, we conclude that systemic lupus erythematosus is potentially represented by a combination of three phenotypes in *C. elegans*: A cell proliferation phenotype (either increased or decreased), probably marked by increased aneuploidy; a developmental phenotype involving cell fate transformation and leading to dysmorphias; and a molecular phenotype involving impairment of the nonsense-mediated decay pathway. The results from the tissue enrichment analysis highlighted three tissues that are particularly sensitive to *lin* mutations (the gonad, the excretory duct cell and the vulval precursor cells), and the gonad arms undergo large quantities of nuclear proliferation.

### Rheumatoid arthritis

Rheumatoid arthritis is an auto-immune disease that is characterized by swollen and painful joints that progressively deteriorate [37]. Unlike lupus, rheumatoid arthritis is not life-threatening, and comorbidity between rheumatoid arthritis and lupus has not been described in past comorbidity studies [38], suggesting that they may have at least partially distinct genetic causes. We found 309 genes associated with rheumatoid arthritis, for which we found 124 worm homolog candidates.

The only phenotype that was enriched for these orthologs was ‘short’ (10 genes, *q* < 10^−4^), even though 64 orthologs were associated with at least one phenotype term. No tissue was enriched in this dataset. Because 82 genes are annotated to have expression in at least one tissue, the lack of enrichment does not reflect ignorance about the sites of expression of these genes. GEA showed that enriched molecular functions for these genes include ‘collagen trimer’ (22 genes, *q* < 10^−15^). However, this term was enriched as the result of degeneracy in the homolog candidates for the SFTPD gene. Other terms included ‘glycosylation’ (10 genes, *q* < 10^−4^) and ‘Golgi apparatus’ (11 genes, *q* < 10^−3^), but these terms were enriched as the result of degenerate homolog candidates for the human gene B3GNT7 which encodes a beta-1,3-N-acetylgalactosaminyltransferase.

The ‘short’ phenotype was the result of the *cat-4*, *dpy-7*, *rnt-1*, *sem-4*, *unc-116*, *ocrl-1* and some genes in the *fat* family. Although these genes are bound by a common phenotype, any genetic relationships between these genes are not immediately clear. Some genes, like *sem-4* and *rnt-1* are likely transcription factors with roles in development (including hypodermal development) [39, 40]. Others are molecular motors (*unc-116*) that are broadly expressed throughout the body of *C. elegans*. Yet others have known roles in neuron and muscle function, such as *ocrl-1* and *cat-4*. The ‘short’ phenotype is a subset of the ‘body length variant’ phenotype. Body length in *C. elegans* can be controlled via cell size or shape [41, 42]; alternatively, cuticle development can alter body shape [43, 44]; finally, muscles can alter the effective body length due to their contraction state [45].

### Obesity-related traits

Obesity-related traits is a category within the GWAS NHGRI-EBI catalog that pools studies that have measured obesity and other traits associated with obesity, such as heart rate, physical activity, hormone levels, body composition and cholesterol levels. Since this category includes many parameters, we expected there would be many phenologs. GWAS studies have identified 957 genes associated with these traits. Using DIOPT, we found 614 orthologs for these genes. In total, 341/614 genes had at least one phenotype annotation; 548/614 had at least one gene ontology term annotation; and 427/614 had at least one tissue term annotation.

Top results for obesity-related traits included ‘acetylcholinesterase inhibitor response variant’ (38 genes, *q* < 10^−6^), ‘neurite morphology variant’ (21 genes, *q* < 10^−2^), and ‘thin’ (31 genes, *q* < 10^−2^). Terms involving locomotion were significantly enriched, as were terms involving body shape and food consumption (*q* < 10^−1^). Concomitant with these phenologs was a tissue enrichment in neuron-related terms. GO enrichment suggested that these genes are participating in ‘iron ion binding’ (40 genes, *q* < 10^−20^) and ‘tetrapyrrole binding’ (37 genes, *q* < 10^−13^).

Tissue and phenotype enrichment therefore suggest that obesity-related traits may be studied in *C. elegans* through neuron physiology and function, specifically with respect to acetylcholinesterase inhibitors. Moreover, GO enrichment implicates iron and tetrapyrrole binding as metabolic components of the obesity-related phenologs in *C. elegans*.

## Ontology enrichment as an aid for screen design

An additional use for a tool like PEA would be as a tool to help guide and design screens to identify genes from an RNA-seq or other genome-wide experiment for further study. This would be particularly useful in cases when researchers may not know what phenotype to expect, in which case PEA can guide selection of a phenotype. Another use case is a scenario where the phenotype under study is not easy to screen for. By finding phenologs to the phenotype of interest, the researcher can design an easier screen for genes that affect the phenolog in question, then re-test genes for the original phenotype of interest.

### Enrichment in the ciliary neuronal transcriptome

As an example of how ontology enrichment can improve our understanding of transcriptomes, we selected a ciliary neuron dataset [19] and ran the complete WormBase Enrichment Suite on it. Ciliary neurons are present in the *C. elegans* male tail, but they are also present in the male cephalic sensillum and hermaphrodites also have ciliated neurons. PEA reveals that the ciliated neuron transcriptome is enriched for genes that are typically associated with ‘meiotic chromosome segregation’ (46 genes, *q* < 10^−5^), ‘aneuploidy’ (42 genes, *q* < 10^−5^) and ‘spindle defective early embryos’ (45 genes, *q* < 10^−2^) (see Fig. 3).

**Figure 3.**
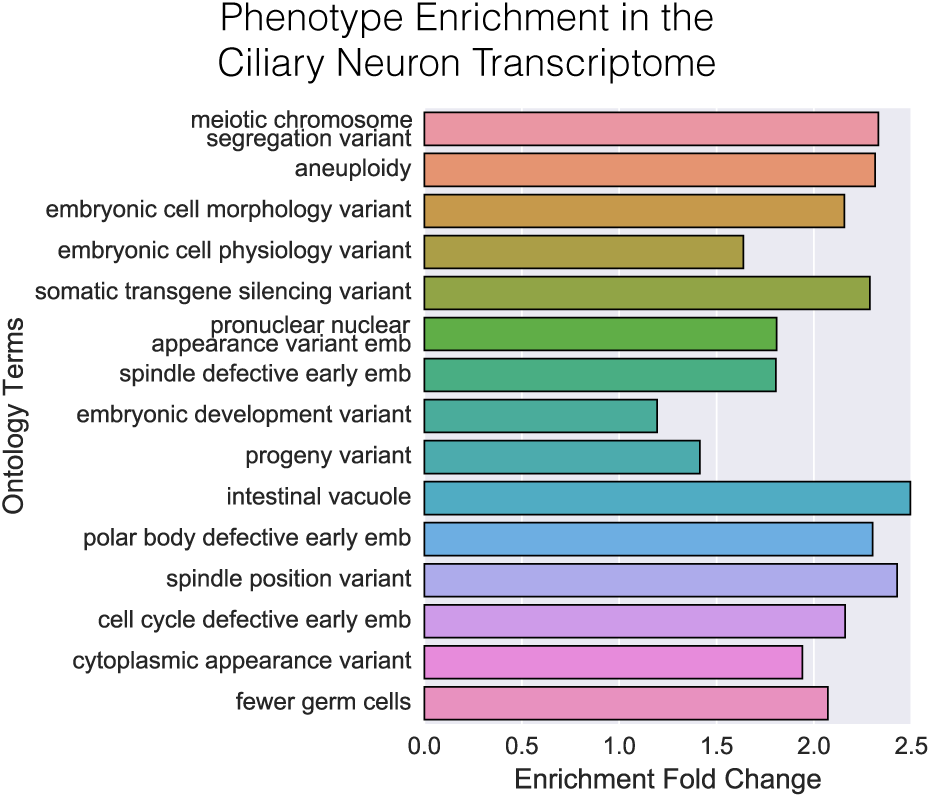
PEA shows that the ciliary transcriptome is enriched in phenotypes related to cell division. Although this could reflect enrichment of microtubules and microtubule-related genes, the enrichment is at least partially driven by cell-cycle and DNA repair genes.

In addition, TEA points at the *C. elegans* gonad primordium, the somatic gonad and early embryonic cells as the sites where genes associated with ciliary neurons are enriched. The ‘male distal tip cell’ is a tissue that is overrepresented in this dataset, but ‘distal tip cell’ is not enriched. In *C. elegans* the hermaphrodite distal tip cells (DTCs) and the male DTC are very similar to each other in their biological functions (both maintain a stem cell niche in the distal gonad). However, the male DTC is non-migratory, whereas the hermaphrodite DTCs are migratory. Therefore, the term ‘male distal tip cell’ may reflect cellular aspects that are correlated with the non-migratory aspects of the DTC biology, such as maintenance of proliferation.

Although one interpretation of the results would be that microtubule genes are driving the enrichment of these terms, another possibility is that there are cell-cycle genes that are driving the enrichment of these phenotypes and tissues. Indeed, GO enrichment shows terms such as ‘DNA replication’ (29 genes, *q* < 10^−5^), and ‘purine NTP-dependent helicase activity’ (15 genes, *q* < 10^−1^). Visual inspection of list in question reveals that cell-cycle and DNA replication/repair genes are abundant in this transcriptome and include genes such as *atm-1*, *dna-2*, or *hpr-17*. This analysis reveals that the ciliary neuron transcriptome is enriched in genes associated with microtubules, but includes genes that are thought to interact with DNA either via repair mechanisms or cell-cycle control [46–48].

## Deconstructing phenotype-tissue relationships

### Tissue enrichment on the Egl gene set reveals cellular components of the phenotype

How does a phenotype emerge? We realized that with the tools that we have developed, it is possible to understand what tissues contribute to a phenotype in a probabilistic framework. In other words, we can extract all genes associated with a particular phenotype, then search for tissue terms that are enriched to understand how a phenotype arises from interactions between anatomical regions. As a test of this, we selected the egg-laying defective (Egl) phenotype. In *C. elegans*, egg-laying is a complex behavior that involves a large number of tissues [49]. The somatic gonad acts as a repository for the eggs, the uterine seam cells help protect the uterus, and a variety of muscles help contract the uterus and open the vulva to lay an egg [50]. The vulva must be well-formed to allow passage of an egg, and the hermaphrodite-specific neuron (HSN) is involved in the egg-laying control [51]. The complexity of the interactions that happen to allow egg-laying make understanding the Egl phenotype in terms of tissues a challenging task.

We extracted all of the *C. elegans* genes that have been associated with an Egl phenotype and we used TEA to understand what tissues are enriched. The HSN was enriched more than five-fold above background (*q* < 10^−7^) as were vulD, vulC, vulE and vulF (*q* < 10^−6^). The vulA, vulB2 and vulB1 were enriched at slightly lower levels (*q* < 10^−5^), whereas the uterine muscles and uterine seam cells were enriched more than twice above background levels (*q* < 10^−2^) (see Fig. 4). Therefore, the Egl phenotype would seem to emerge primarily from defects in the HSN, secondarily from defects in the vulva, and only sometimes from defects in the uterine seam cells or muscles. It is notable that all vulval cells were not equally enriched. Although all the ‘vul’ cells are annotated to a similar degree (between 50–70 genes for each cell type), the vulD and vulC cells had the largest enrichment effect size and the lowest q-values, suggesting that these cells are more likely to be associated with an Egl phenotype than the others. This may reflect the fact that vulD and vulE are the site of attachment for four vulval muscles, vm1. Perhaps this attachment is particularly fragile, and perturbations to these cells prevent adequate function of these muscles. In support of these observations, P7.pa had the largest fold-enrichment of any tissue. In *C. elegans*, P7.pa gives rise to vulD and vulC. However, vulF is also attached to a set of four additional vulval muscles, vm2. Why is vulF less associated with an Egl phenotype?

**Figure 4.**
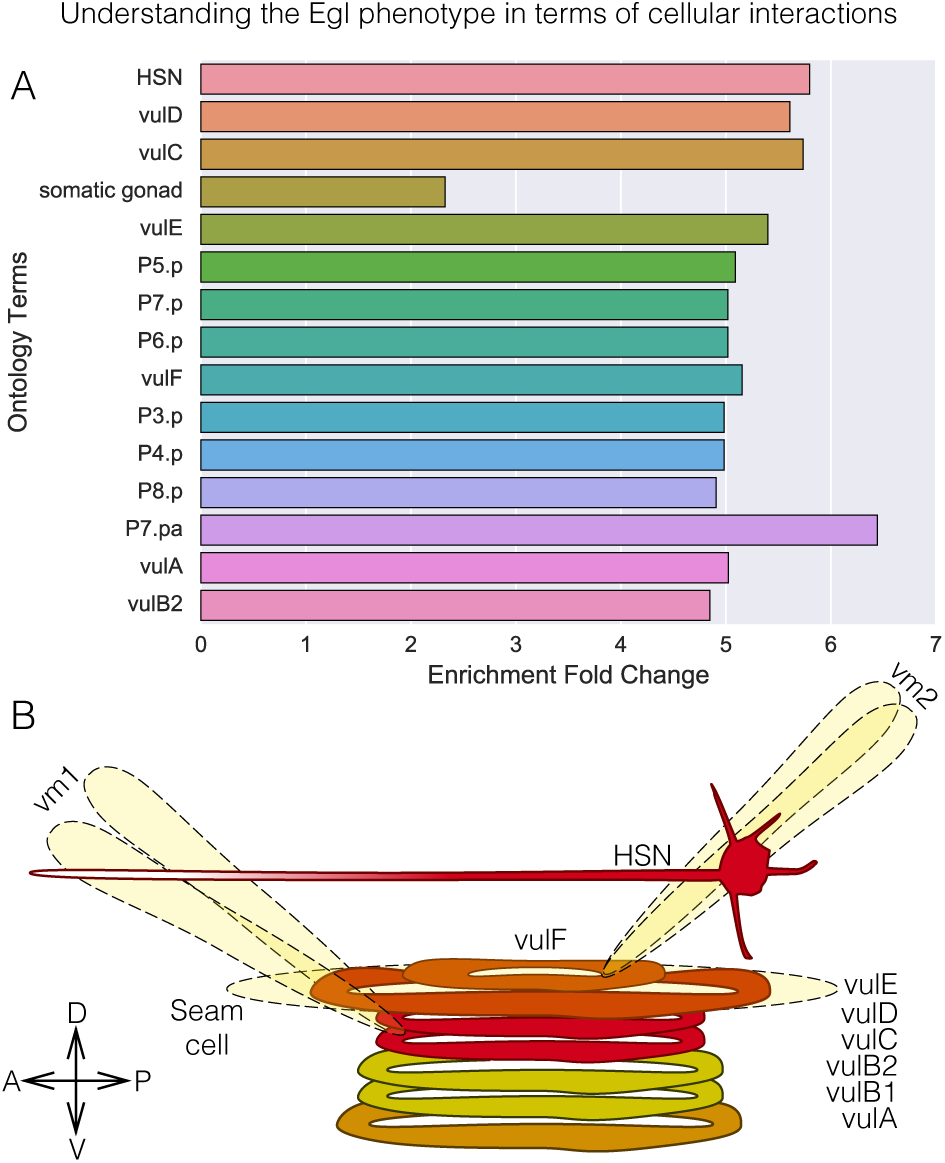
The Egl phenotype is a complex phenotype that is the result of interactions between many tissues. To dissect the contributions of individual tissues to generating an Egl phenotype, we obtained all genes annotated with with an Egl term. **A** We used TEA on the set of Egl genes to identify enriched tissues. **B** Anatomic diagram showing the tissues that are most enriched in the Egl gene set. Color coding shows the qualitative ordering of enrichment (red-Most enriched, yellow-least enriched). For clarity, not all cells are shown. All vm1 (4 cells) and vm2 (4 cells) muscles are symmetrical arranged around the vulva, but only 2 cells are shown for each. There are two seam cells on the left and right side of the vulva, but only cell on the right is shown. Only terminally differentiated cells are shown.

### Quantifying the anatomy-phenotype mapping via Bayesian probabilities

Another way to understand the phenotype-anatomy mapping is by considering how informative a given anatomy term is on a particular phenotype, or vice-versa. To this end, we calculated two conditional probabilities that helped us answer this question. The first conditional probability,

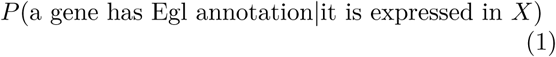

answers the question: For a gene with an expression pattern that includes the tissue term X (i.e., the gene is expressed at least in X), what is the probability that this gene has an Egl phenotype (i.e., the phenotype annotations for this gene include Egl)? For simplicity, we can rewrite this equation more succintly by removing a few words. The calculation of this probability is straightforward and follows from the definition of conditional probability:

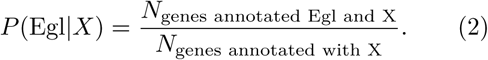

Equation 2 measures how likely a gene is to be annotated with an Egl phenotype given that its expression pattern includes the term *X*. A related quantity (which is neither the inverse nor the complement) is the conditional probability that a gene which is annotated with at least the Egl phenotype is expressed in tissue *X*. That is to say,

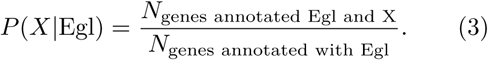

Equation 3 tells us how probable it is that any given gene that is annotated with an Egl phenotype includes *X* as a tissue term. Taken together, equations 2 and 3 help us understand how predictive anatomic expression is of phenotypes, and how predictive phenotypes are of anatomic expression.

We calculated the conditional probability that a gene has an Egl phenotype given that it’s expression pattern includes a tissue term *X* and we searched for the tissue terms that maximized this probability. The list of terms that maximized this probability reflected the results from running TEA on the subset of genes that have an Egl phenotype. We also calculated the conditional probability that a gene has expression in a tissue term *X* given that it is annotated with an Egl phenotype and we searched for terms that maximized this probability (see Table 1). The terms that maximized this probability were ‘nervous system’, ‘pharynx’ (a body part with a lot of neurons), ‘sex organ’ and ‘tail’ (a body part with neurons and hypodermis). In general, the terms that had a high *P*(Egl|*X*) did not have a high *P*(*X*|Egl). Additionally, the terms that had a high *P*(*X*|Egl) are broad terms that include a lot of cells, whereas the terms that had a high *P*(Egl|*X*) were considerably more specific. We conclude that the Egl phenotype arises from a small set of tissues. The Egl phenotype can be best predicted by genes with expression patterns that include at least one of a small number of cells (mainly vul cells, HSN). On the other hand, answering whether the expression pattern of a gene includes a particular anatomic region or tissue given that the gene has an Egl phenotype is hard to do for small tissues or single cells. However, guesses about what functional system or broad anatomic region an Egl gene is expressed in can be answered with confidence (~ 70% of the time, an Egl mutant is expressed in the nervous system).

**Table 1.** Conditional probabilities for various tissues. The first column shows the conditional probability that agene has an Egl phenotype given that it has expression in tissue *X* (given by the row). The second column shows the conditional probability that a gene has expression in the anatomy term X given that it has an Egl phenotype. The first 9 terms are the terms for which *P* (Egl|*X*) is maximized. The last three terms are the terms which have the highest *P*(*X*|Egl). For clarity, the Pn.p cells are not shown even though *P*(Egl|Pn.p) ~ 0.24.

| Tissue | $P(\text{Egl}\vert X)$ | $P(X\vert \text{Egl})$ |
| --- | --- | --- |
| P7.pa | 0.30 | 0.04 |
| HSN | 0.27 | 0.11 |
| vulC | 0.27 | 0.06 |
| vulD | 0.26 | 0.07 |
| vulE | 0.25 | 0.06 |
| vulF | 0.24 | 0.06 |
| vulA | 0.24 | 0.05 |
| vulB2 | 0.23 | 0.05 |
| vulB1 | 0.22 | 0.05 |
| nervous system | 0.02 | 0.72 |
| pharynx | 0.00 | 0.46 |
| sex organ | 0.07 | 0.41 |
| tail | 0.05 | 0.33 |

## Conclusions

The addition of GO and Phenotype Ontology enrichment testing to WormBase marks an important step towards a unified set of analyses that can help researchers to understand genomic datasets. These enrichment analyses will allow the community to fully benefit from the data curation ongoing at Worm-Base. By using the same algorithms to generate enrichment dictionaries for testing and using the same model to test for term enrichment, these tools provide a coherent framework with which to analyze genomic data. In particular, it is our hope that phenotype enrichment will be of use to geneticists performing genome-wide analyses, because they are familiar with the ontological terms that are tested and with their biological meaning. Ideally, PEA could allow researchers to design better screens to maximize target gene identification by quantifying the phenotypes that are most overrepresented. An intriguing new direction of research would be to create a controlled language for screening methods. With such a language, we should be able to computationally suggest to a researcher what screens one may wish to perform given a dataset.

